# Phylodynamics without trees: estimating R0 directly from pathogen sequences

**DOI:** 10.1101/102061

**Authors:** Giacomo Plazzotta, Caroline Colijn

## Abstract

We develop a new tree-free phylodynamic method to estimate the reproduction number (R0) of a pathogen from large numbers of sequences of a pathogen. It is based on the convergence of the cherry-to-tip ratio (CTR) to a constant depending on R0 in supercritical branching trees. It is a tree-free method because tree reconstruction is not required: the number of cherries and the CTR is estimated directly from the sequences using the new computational method Cherries Without Tree (CWT). With simulations, we compare CWT to other methods currently in use. We use the new inference method to estimate R0 from simulated sequences and discuss its accuracy. We explore the potential bias arising from sub-sampling.

## 1 Introduction

Phylogenetic trees derived from pathogen sequence data are used to describe the spreading patterns of the pathogen, in a growing field known as phylodynamics [1, 2]. The underlying assumption behind phylodynamics is that genetic variability contains information about the population’s recent ancestry and demographics, and in the case of pathogens, this information can help to help curtail the spread of infectious disease.

The basic reproduction number *R*_0_ is a fundamental epidemiological parameter: the average number of secondary infections an infectious individual causes over the infectious period, in a fully susceptible population. Estimating the basic reproductive number can be challenging, and typically requires simplifying assumptions together with knowledge of the duration of infectiousness, the epidemic growth curve, the number of susceptibles at equilibrium, the average age at infection at equilibrium, or a closed population [3]. This is particularly difficult for pathogens whose infectious
period is long and variable, and/or for which case identification is challenging.

There are several approaches to estimating *R*_0_ using genetic data, largely based on Bayesian inference. These include inferring the parameters of a birth-death process [4, 5, 6], coalescent methods based on Bayesian skyrides or epidemic coalescents, [7, 8] and fitting epidemiological models to phylogenetic trees to estimate epidemiological parameters [9, 10, 11]. Indeed, part of the motivation for phylodynamics is that there are important pathogens for which we have the opportunity to gather large numbers of genetic sequences but we do not have reliable estimates of epidemic parameters such as *R*_0_; these range from the common cold to tuberculosis. In practice, as real populations are rarely fully susceptible, methods based on genetic data estimate an effective reproduction number *R* rather than strictly *R*_0_.

To date, all approaches are computationally intensive and are unlikely, even with advanced MCMC techniques, to scale well to large datasets. This is exacerbated by the fact that most approaches either require or infer a timed phylogenetic tree, a task that in itself is very challenging for large numbers of sequences. Furthermore, if underlying parameters are the central quantity of interest, proceeding via phylogenetic trees or transmission trees is not always desirable, as the number of possible trees and the difficulty of computing relevant likelihoods is a barrier.

Here we present a tree-free way to estimate *R* from large numbers of sequences. It is based on the fact that the number of cherries is closely linked to the basic reproduction number [12]. The number of cherries is a simple shape statistic that does not depend on deep branching, timing or other complex questions in phylogenetic inference. It can be used to infer *R*_0_ from sequence data where there are large numbers of sequences. In addition, it is not necessary to reconstruct the whole tree to obtain the number of cherries and to infer *R*_0_. Indeed, tree reconstruction methods carry underlying modelling assumptions and could cause "shape bias" [13]. Our technique estimates the number of cherries directly from the genetic data, without tree reconstruction, and infers *R*_0_ from the numbers of cherries. (If sequences are derived from populations that are partially immune the method infers *R*). We test the approach on simulated trees and sequences, apply it to estimate the reproduction number of the 2009 H1N1 epidemic and of *S. typhi*, and explore the robustness of our approach to partial sampling.

## 2 Methods

### 2.1 Cherries and *R*_0_

Many approaches in phylodynamics work from the assumption that information about transmission is encoded in the branching events in phylogenetic trees. When hosts carry limited pathogen genetic diversity, branching events in timed phylogenetic trees can be used as a model for transmission events in the host population (see for example [14, 2, 15]). Even when there is pathogen diversity within hosts, it is often the case that inferred transmission events are close to phylogenetic branching events [16]. Our approach uses the simplifying assumption that branching events in the outbreak process correspond to branching events in the genealogical tree of the pathogen.

The ratio between the number of cherries and the number of tips converges in distribution to a single number, depending on *R*_0_, as the number of tips approaches infinity; the form of this relationship depends on how infectivity changes over the course of infection. The link between the basic reproduction number and phylogenetic or genomic data is found using Crump-Mode-Jagers branching processes to model outbreak trees as in [12]. In this setting, each tip corresponds to an individual’s death/removal and each internal node corresponds to a birth/infection event. A cherry is a configuration made of two tips joined together [12, 17, 18]. We recently found a close link between the number of cherries in branching trees and *R*_0_ [12]; in Crump-Mode-Jagers branching processes, the cherry-to-tip ratio (*CTR*) converges almost surely to a constant as the tree grows. This constant depends on *R*_0_,or *R* in the case that depletion of susceptibles affects the offspring distribution. (For the remainder of this work we will refer to *R*_0_, with the understanding that the method estimates the mean of the offspring distribution in an underlying branching process, equivalent to either *R*_0_ or *R* depending on whether depletion of susceptibles has occurred in the relevant populations; this caveat applies to other sequence-based phylodynamic methods).

In the homogeneous case, where births and deaths occur at constant rates, the cherry-to-tip ratio reduces to 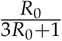 for large trees. Here, we use this relationship to develop a method to estimate *R*_0_. The approach is summarised in Figure 1. The point estimate for *R*_0_ is:

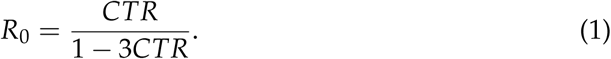

**Figure 1:**
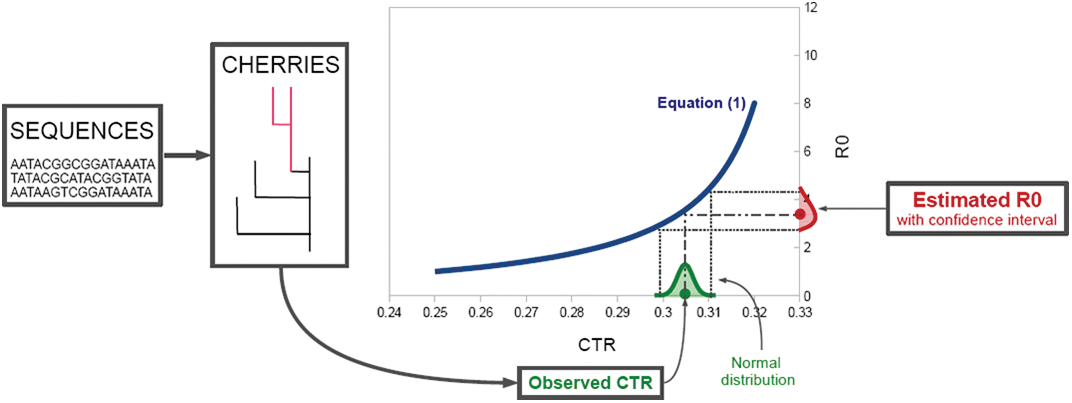
Conceptual figure describing the steps of the *R*_0_ inference. A cherry is any sub-tree with two tips (illustrated in pink).

In the constant rate model, infections/births and removals/deaths both occur at a constant rate, but this can be generalised using knowledge of the relationship between the number of cherries in systems with non-constant rates and *R*_0_ [12]; the specific form of the relationship between *R*_0_ and the CTR depends on the natural history of infection.

The distribution of the CTR around its limit converges to a normal distribution (Supporting Information), and the variance decreases, approaching 0 as the number of tips grows. We use the limiting normal distribution to find error bound on the inference of *R*_0_ using the numbers of cherries:

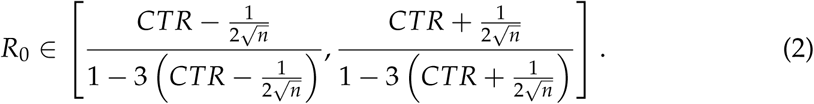

### 2.2 Simulations

#### Testing the Cherries Without Trees algorithm

The tree simulations were carried out in R [19] with function pbtree of package phytools [20]; the parameter n was set at 150, 700, 1675, to provide trees with 500, 2000, and 5000 tips on average respectively. The parameters b and d for birth and death rates were 1.5 and 1 respectively, corresponding to an *R*_0_ value of 1.5. For each group, 25 trees were simulated. The simulated sequences were obtained with the function SimSeq from the R package phangorn [21] and had 20000 characters, the rate parameter was 0.03. These simulations underly Figures 2 and 3.

#### Testing the confidence interval bounds

(Section 3.3). The tree simulations were carried out similarly as in the previous paragraph: in R [19] with function pbtree of package phytools [20], but with the parameter n was set at 700, 1675, and 3350 so that, on average, trees had 2000, 5000, and 10000 tips respectively. The parameters b and d for birth and death rates were 1.5 and 1 respectively, corresponding to an *R*_0_ value of 1.5. For each group, 25 trees were simulated. Sequences were simulated using the function SimSeq from the R package phangorn [21] and had 20000 characters, the rate parameter was 0.03. The algorithm CWT was used to estimate the number of cherries on these sequences and provide *R*_0_ estimates.

#### Testing the effects of sampling on the CTR

Section 3.5). Using a modified version of function pbtree [19, 20] built for this purpose, we simulated three large trees (about half a million tips) and *R*_0_ 1.5, 2, and 2.5 respectively. The modification of pbtree was necessary to speed up the computing time, and consists two phases: simulation of a first tree with *n* currently alive tips, and simulation of further *n* trees cut off at a predefined time *t*. Function drop.tip [19, 22] was used to perform independent prunings of the trees at different sampling levels (ten pruning per level).

## 3 Results

### 3.1 *R*_0_ from the cherry-to-tip ratio in reconstructed trees

The cherry-to-tip ratio converges to the value in Eq. 1, as the trees used grow larger (see Figure S1), following the theoretical result that the variance decreases to 0. Inference approaches based on the number of cherries are therefore applicable to large datasets with thousands of sequences, whereas established phylodynamic approaches are well-suited to datasets with at most hundreds of sequences.

We considered a publicly available tree with 51169 tips derived from *Salmonella typhi* sequences [23]. The number of cherries of the whole tree is 15323, corresponding to an *R*_0_ estimate of 2.95, with bounds [2.57, 3.44] from Eq. (2). This is consistent with other studies: an *R*_0_ of 2.8 was estimated in Vellore (India) [24]; a value of 3.4 was found for the reproduction number from an untreated clinical infection in [25]. However in Dhaka, Bangladesh, *R*_0_ for typhoid fever was estimated at 7 [24] (but not all cases of *S. typhi* cause typhoid fever).

The reconstructed tree contains unnecessary information for the *R*_0_ estimation using equations (1) and (2) because these relationships only require the number of cherries. We developed a technique that can estimate this number directly from sequence data.

### 3.2 A tree-free technique to estimate cherries

Although the formulas and derivations of this paper are based on phylogenetic tree analysis, the central formula of equation (1) only requires the *CTR* for the estimation of *R*_0_. This is an alternative starting point for Equation (1) to infer *R*_0_. We have developed an algorithm to find the cherry-to-tip ratio without first inferring a phylogenetic tree, and we refer to the technique as CWT for *Cherries Without Trees*.

Because the choice of the root itself often carries uncertainty while adding/deleting at most one cherry, we navigate the sequences in an unrooted manner. First, we consider a tip *c* and seek the tip, if any, that forms a cherry with *c*. CWT is based on a quartet selection. Given any four sequences, there are only three unrooted quartets whose tips correspond to the given sequences. We choose one of these quartets as the “right” one, with a consistent method previously determined (parsimony, distance methods, maximum likelihood, etc). Once the quartet is selected, only one tip among the three could potentially form a cherry with *c* in the whole tree, namely the unique tip in the best quartet that is in a cherry with *c*; we name it *m*. The other two sequences cannot form a cherry with *c* (because *m* is a better candidate), and we name one of them *f* (for "false") and discard the other. This is the initialisation phase, where we identified a potential candidate *m*, and a particular wrong candidate *f*. The algorithm then applies the quartet selection systematically to all the remaining tips and updates *m*, *f* and *t* (refer to the pseudo-code for more details). Those sequences for which there is an *m* that is a good cherry candidate at the end of this testing are tips that are in an inferred cherry. Importantly, the accuracy of CWT depends on the accuracy of the quartet selection mechanism: if the quartet selection always returns the right quartet, then CWT returns the exact number of cherries. The quartet is the minimal configuration to investigate because with three unrooted tips, a cherry is not well defined.

Note that the candidate *m* can have status *good* or *bad*. If the quartet at the bottom in Table 1 is chosen, neither *m*, *f*, or *t* can form a cherry with *c* in the whole tree. But *m* is kept as a candidate with status *bad* to inform new iterations of the algorithm. It is not redundant to loop *t* among all the sequences, even if some of them are known to form other cherries (saving information from previous loops). If we discard the known cherries from the loop, new cherries would be falsely detected. For instance consider a caterpillar tree, i.e. a tree with a single cherry, and delete that cherry. A new one would automatically appear, and so on.

**Table 1:**
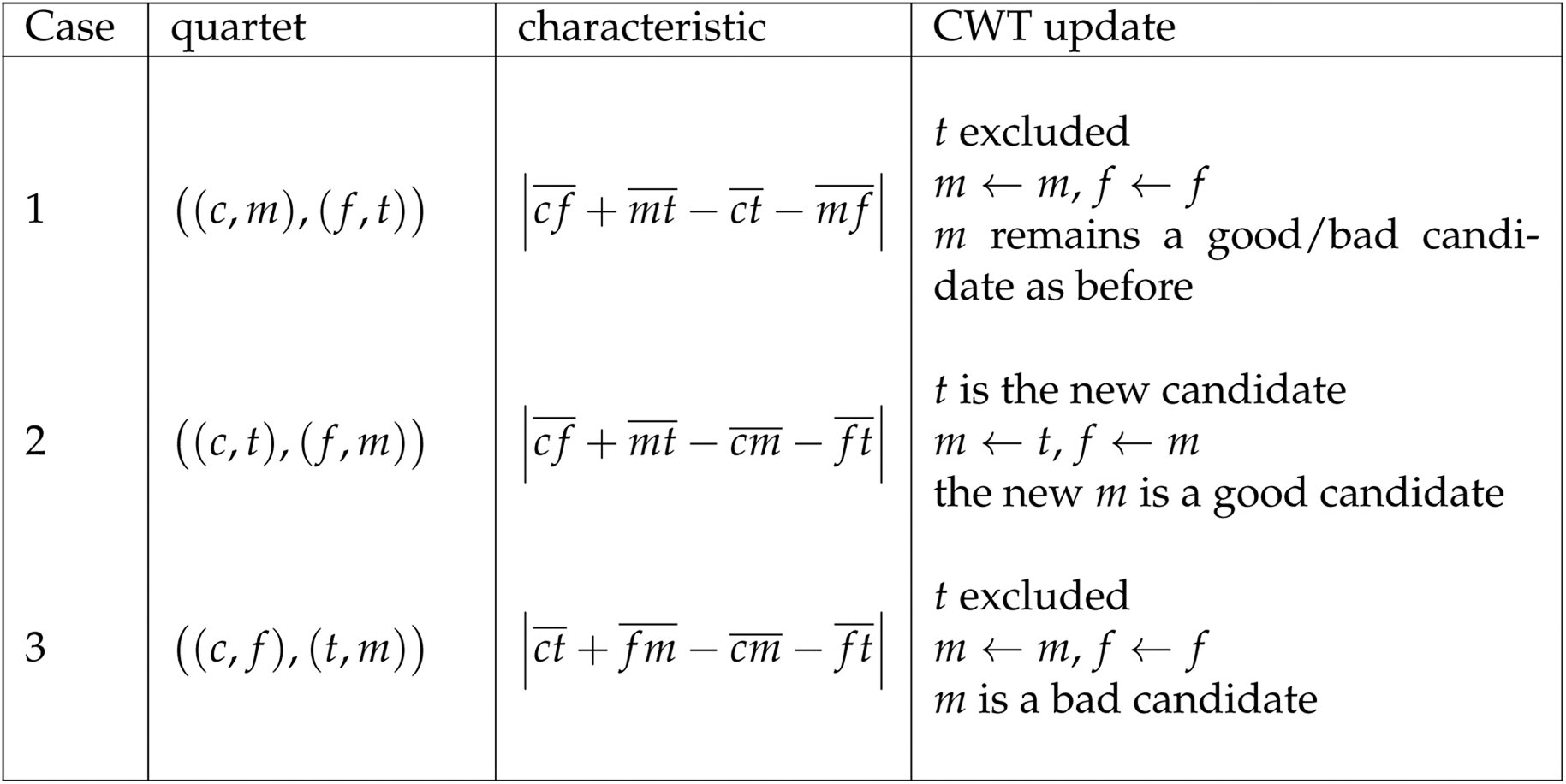
Quartet selection performed in the tree-free algorithm to estimate the number of cherries from sequences. The labels *c*, *m*, *f*, and *t* correspond to the sequences of the quartet. The over line symbol 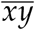 correspond to the genetic distance between *x* and *y*. If the distance measure considered is “tree like”, then only one of the characteristics equals zero, and it corresponds to the correct case. However genetic distances are not tree like in general, and none of the characteristic is exactly zero. Here we use the Jukes Cantor distance [26] and pick the case corresponding to the minimum. If the minimum is larger than 0.2 times any other, the `flag` in the algorithm is set to 1. This occurs in quartets in which the internal branch length is much shorter than the 4 pendant lengths, and is related to long branch attraction. To address this we resample *f* (from the previously-tested *f* nodes, then previously-excluded *m*’s and *t*’s, because we know that these do not form a cherry with *c*) until the three cases can be distinguished. We chose the 0.2 threshold empirically based on performance.

The accuracy of CWT cannot be tested on real genetic data because the true number of cherries is unknown; we tested the approach using simulated sequences evolved on known trees (with known *R*_0_; see Methods). We ran CWT as well as inferring trees using neighbour-joining (NJ) [27] and FastTree [28, 29], and Figure 2 illustrates the results. We used a t test to compare the relative error in CTR values to the null hypothesis of zero bias (ie mean relative error equals zero). The CWT algorithm slightly (and significantly) underestimates the cherry-to-tip ratio, with a mean relative error of -0.011, 95% confidence interval at [-0.019, -0.003], and a p-value of 0.0051 in the t-test. NJ consistently overestimates the CTR with a relative error of 0.221, 95% confidence interval at [0.214,0.228], and a p-value < 10^−15^ indicating very significant departure from a mean of 0. `FastTree` also overestimates the CTR, with relative error 0.058, a confidence interval of [0.053, 0.062] and a p-value < 10^−15^ (also significantly away from 0). While CWT showed significant bias, the bias is small and the p-value (0.0051) is comparatively high. Figure 2 compares the estimated and true CTR values for neighbour-joining, `FastTree` and our CWT algorithm and illustrates that CWT is on average comparable to FastTree and much more accurate than neighbour joining in estimating the number of cherries.

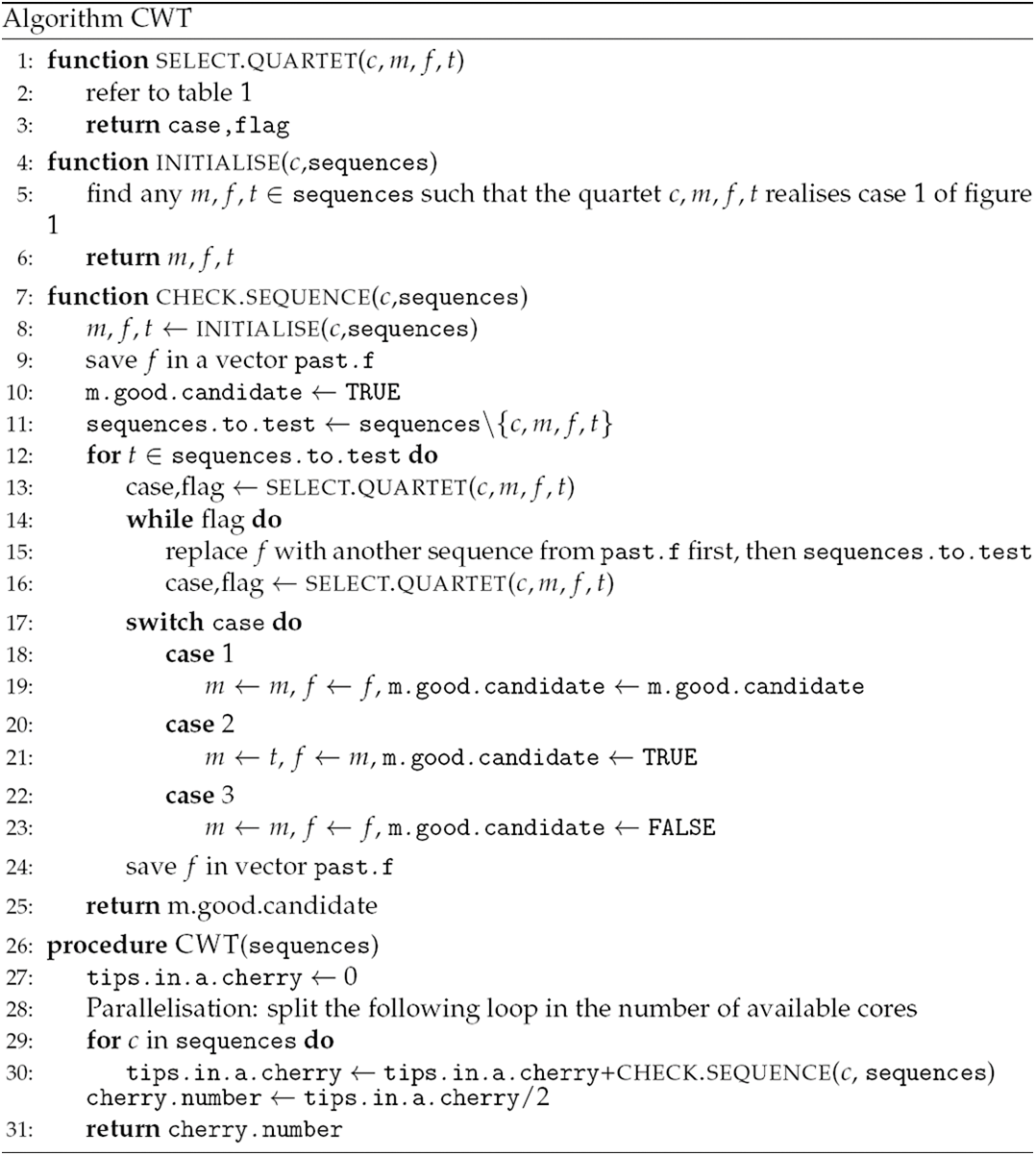

**Figure 2:**
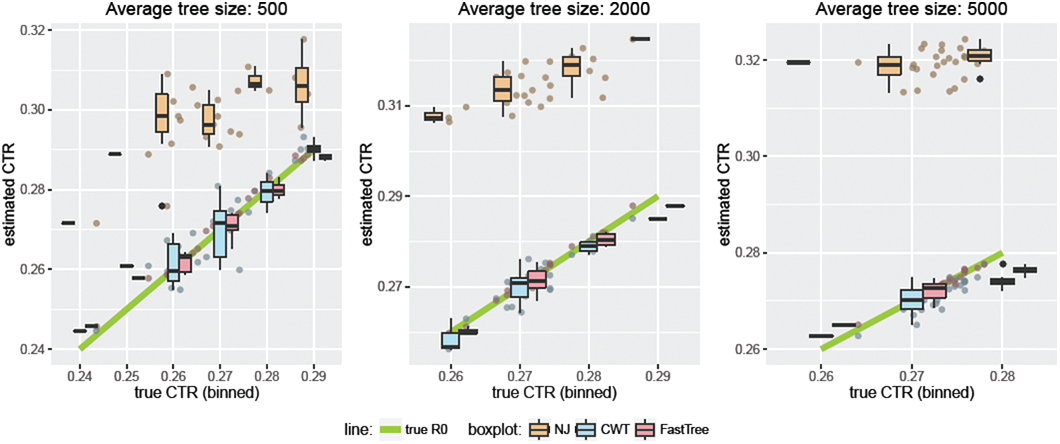
The estimated CTR using the algorithm CWT (blue) is compared to the real CTR (green line), the estimation using NJ (orange), and the estimation using `FastTree` (red). The tree simulation were carried out in `R` [19] with function pbtree of package phytools [20]; the parameter n was set at 150, 700, 1675, to provide trees with 500, 2000, and 5000 tips on average respectively. The parameters `b` and `d` for birth and death rates were 1.5 and 1 respectively, corresponding to an *R*_0_ value of 1.5. For each group, 25 trees were simulated. The simulated sequences were obtained with the function `SimSeq` from the R package `phangorn` [21] and had 20000 characters, the `rate` parameter was 0.03.

For each of the *n* tips, CWT loops through other *n* − 3 tips doing quartet selection. The time of quartet selection does not depend on *n*, in fact it requires *O*(*l*) operations using the Jukes/Cantor distance, where *l* is the sequence length. Therefore CWT before parallelisation has a time complexity of *O*(*ln*^2^). CWT can be parallelised up to *n* times because each iteration of the wider loop is independent. This would reduce CWT’s time complexity to *O*(*ln*). In terms of memory, each iteration needs to store only a few scalars represending tip indices, so the memory required does not depend on *n* and is *O*(1) or *O*(*l*) depending on the genetic distance function chosen. In comparison to tree inference, just the computation of the distance matrix between sequences has a time complexity of *O*(*ln*^2^) and requires *O*(*n*^2^) memory space, setting a minimum threshold for all distance matrix methods like NJ. NJ can be performed with extreme efficiency [30], reducing its time to the computation of the distance matrix itself. On the other hand, FastTree is better, requiring just 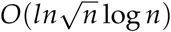 time and 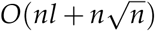 memory [28]. CWT outperforms any other methods in terms of memory (though of course it only computes the number of cherries). If parallelised enough times, CWT could also outperform other methods in terms of time. On a standard desktop with Intel^®^ Core^™^i7-3770S CPU, for 1000 sequences of length 20000 FastTree took 675 seconds, and CWT took 253 seconds (parallelised in 7 nodes). We estimate that CWT would take 4-8 seconds if parallelised 1000 times. The current version of CWT is built within R and can be optimised even further if written in lower-level languages such as C++.

### 3.3 Tree-free inference of *R*_0_

We simulated sequences evolving in a process with a known *R*_0_ (1.5), used CWT to estimate the number of cherries, and then inferred *R*_0_ from the cherry-to-tip ratio. Figure 3 illustrates the results, and highlights the convergence as the number of tips grows. We computed the confidence interval (Equation 2), and found that the true *R*_0_ was outside the confidence interval only 121 times over 5000 trees (see *Testing the confidence interval* in the methods section), corresponding to a level of confidence of over 97.5%. The *R*_0_ point estimate from equation (1) had an average relative error of 2.75%. It is a challenge to analytically derive the level of confidence for the interval in equation (2). However, from equation (eq:varUp), 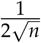 is an upper bound for the standard deviation of the *CTR*, and it therefore guarantees that the interval in equation (2) has a confidence of at least 70%, while our empirical results suggest that the confidence is much higher.

**Figure 3:**
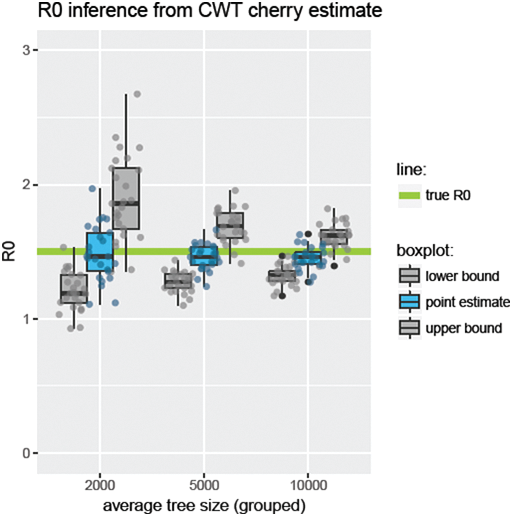
The *R*_0_ point estimate from equation (1) and upper and lower bounds from equation (2). The simulated trees and sequences are the same used for figure 2. The number of cherries was estimated directly from the sequences with algorithm CWT. The precision of this method of inferring *R*_0_ increases as the number of sequences considered increases. The tree simulation was carried out in R [19] with function pbtree of package phytools [20]; the parameter n was set at 700, 1675, 3350 to provide trees with 2000, 5000 and 10000 tips on average respectively. The parameters b and d for birth and death rates were 1.5 and 1 respectively, corresponding to an *R*_0_ value of 1.5. For each group, 25 trees were simulated.

### 3.4 Tree-free estimate of the reproduction number of H1N1

We downloaded 2975 H1N1 influenza sequences from the *NCBI influenza virus resource* [31] detected from April 2009 onward to September 2010, corresponding to the 2009 pandemic [32]; the accession numbers can be found in the supplementary material. The sequences were aligned using MAFFT [33, 34]. The algorithm CWT was used to estimate the number of cherries directly from the sequences, then equations (1) and (2) were used to provide the estimate and bounds of *R*_0_. Another estimated number of cherries was derived from the tree reconstructed with FastTree [28, 29], and the resulting *R*_0_ estimates are compared in Table 2. The *CTR* derived with CWT is 0.2704, and corresponds to an *R*_0_ estimate of 1.43 (1.21-1.73) which is in line with other estimates of the 2009 H1N1 influenza pandemic [8, 35, 36]. The *CTR* from the reconstructed tree with FastTree is higher at 0.28303, realizing an *R*_0_ estimate of 1.88 (1.53-2.36), which is slightly higher than most estimates but still within most margins of errors [36].

**Table 2:** Estimation of *R*_0_ of H1N1 influenza during the 2009 pandemic using equations (1) and (2). The *CTR* was calculated directly from 2975 sequences using CWT, or from the the tree reconstructed with FastTree [28, 29]. The *R*_0_ estimates and the bounds are calculated from the *CTR* with equations (1) and (2) respectively. The 2975 sequences were downloaded from the *NCBI influenza virus resource* [31], and correspond to all full non-identical sequences sampled between April 2009 to September 2010. The accession number can be found in the supplementary material.

| Method | CTR estimate | $R_0$ estimate (upper bound - lower bound) |
| --- | --- | --- |
| CWT | 0.27042 | 1.43 (1.21 - 1.73) |
| FastTree | 0.28303 | 1.88 (1.53 - 2.36) |

### 3.5 Effects of sampling on the the *CTR* and *R*_0_ point estimate

The convergence results in [12], and so equation (1), are in the context of fully sampled trees. However, data on infectious disease outbreaks typically would have at least a few missing cases due to the challenges of ensuring that every case is identified, sampled, sequenced and included in a dataset. While the *CTR* would converge in any branching process, fully or partially sampled, the limit for a sub-sampled process may differ from the expression in Equation (1). Deriving analytic convergence results for the *CTR* that take sampling into account is challenging because of the variability of the processes involved (removal and reconstruction of the pruned tree). We explore sub-sampling large phylogenetic trees to model the effects of incomplete sampling. To mimic incomplete data and determine how the *R*_0_ estimate is affected, we randomly (uniformly) remove a specified fraction of the tips/nodes of a complete tree and reconstruct a new “sampled” tree.

We found that random uniform sub-sampling has some effect on the *CTR* and on *R*_0_ inference. We simulated three large trees (about half a million tips), each corresponding to one value of *R*_0_: 1.5, 2, 2.5. Using the R function drop.tip (in ape [22, 19]) we randomly and independently pruned tips at different sampling rates. Ten independent prunings at each sampling level were done to avoid outliers. We computed the *CTR* and the *R*_0_ estimate using equation (1) for each pruned tree. Figure 4 shows how the *CTR* varies and related *R*_0_ estimate vary as a consequence of sampling. Close to 0% sampling (almost all tips pruned) the *CTR* converges to 1/3, which is the *CTR* of the only tree topology with three tips. Because the trees are very large, at 100% sampling (no pruning) the *CTR* observed is very close to the limit *R*_0_/(3*R*_0_ + 1) discussed in section 2.1 and related to equation 1. If the sampling rate is lower than 20%, the *R*_0_ estimate can be significantly higher than the true *R*_0_.

**Figure 4:**
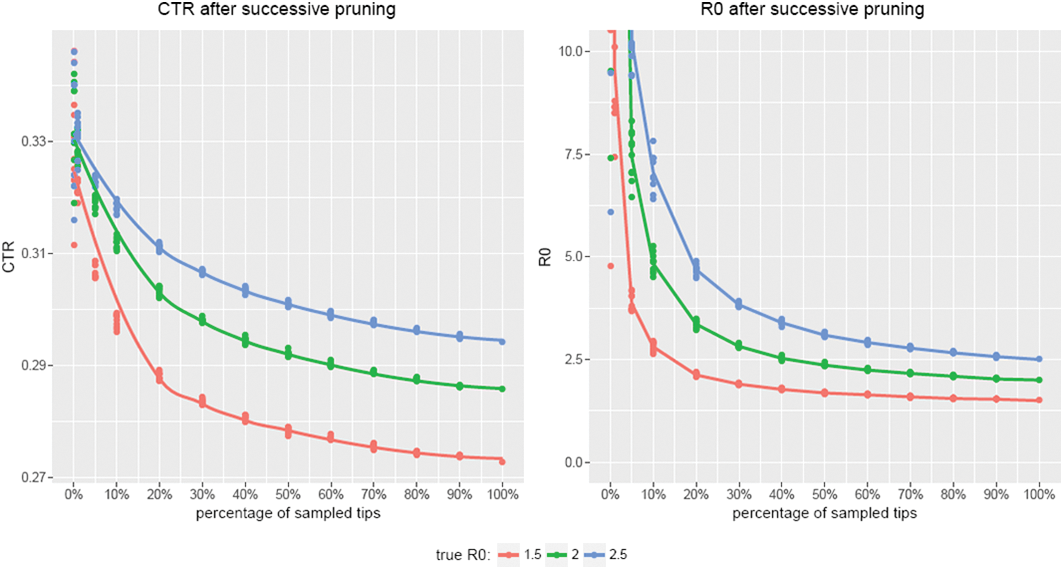
*CTR* computation and *R*_0_ estimation using equation (1) of repeated random pruning of large (≈500K tips) simulated trees. Each line corresponds to a different tree, that was simulated with a modified version of `pbtree` (`R` package `phytools` [20, 19]) that allowed for large tree simulation. Each tree corresponds to a different pre-set theoretical value of *R*_0_: 1.5, 2, and 2.5. At each chosen sampling level, a portion of tips were chosen randomly and deleted with a modified version of `drop.tip` (`R` package `ape` [22, 19]). This was repeated ten times for each sampling level, to provide a more robust estimate.

## 4 Discussion

The method we have presented allows *R*_0_ estimation directly from a set of pathogen sequences; it is based on a tree-free estimate of the number of cherries in the true phylogeny of the sequences, and a confidence interval for *R*_0_ based on the cherry-to-tip ratio (*CTR*) [12]. The cherry-to-tip ratio can also be computed from a reconstructed tree simply by counting the number of cherries, providing a tree-dependent estimate of *R*_0_. The accuracy of the method increases considerably when a large number of sequences is available.

We developed the CWT as an attempt to disentangle *R*_0_ estimation from the challenging problem of tree reconstruction. The nucleus of the algorithm is a quartet selection, i.e. the identification of the quartet subtree that links any four tips. If the quartet selection mechanism is exact, then CWT has no errors. Unfortunately, we do not have an exact quartet selection, and have used a distance-based condition. Where a quartet has short internal branch lengths, the noisiness of phylogenetic distance means that CWT is subject to error (analogous to long branch attraction), which we have addressed by allowing resampling of tip *f* in ambiguous quartets.

There are several limitations. By its nature, CWT (or the related tree-dependent *R*_0_ estimate from the number of cherries in the tree) require very large numbers of cases to produce tight estimates. It might be possible to improve on the upper bound that we obtained for the variance, particularly for homogeneous processes, but the required calculations are extremely complex. In using equation (1) we also implicitly assume that the *CTR* is an unbiased estimator of its limit (which is consistent with simulations). For large trees, any bias would be insignificant compared to the large margin of error provided by this bound of the variance, but further investigation may tighten the bound. That said, sequencing technology is becoming widely used in epidemiology; studies with thousands of bacterial genomes are published and researchers are moving to tens of thousands; the CRyPTIC project is sequencing 100,000 TB genomes [37]. Viral genomes are already available in very large numbers. Our approach is relevant to large datasets where conventional approaches are infeasible.

We explored robustness to uniform sub-sampling; this appears to bias to a larger estimate significantly when the sampling rate is lower than 20%. However, non-uniform sub-sampling is likely to be commonplace, and be characterised by a collection of highly sampled subgroups of the population. This would likely reduce the bias as compared to uniform sampling (which is a worst case scenario in this context). If uniform sampling is in fact a good assumption, figure 4 suggests that for a set value of *R*_0_, the curves relating the *R*_0_ estimates to the sampling rates do not intersect. Therefore, given the estimated *R*_0_ and the sampling rate, it is possible to estimate the true *R*_0_.

Despite these limitations, there are clear advantages to our approach. It is tree-free and so avoids the need for expensive tree reconstruction, inference of a timed phylogenetic tree and subsequent MCMC analysis on this tree. It is therefore quick and highly parallelisable, and is suited to very large datasets. Its *O*(*ln*^2^) complexity compares well to FastTree’s 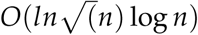 because it is so parallisable, and its memory requirements are much less. Furthermore, it is the basis for *alignment*-free approaches to estimating *R*_0_, because all it requires is an alignment-free distance function (such as a k-mer distance) with which to compare quartets. It is therefore feasible for recombining pathogens or other systems for which alignments are challenging to create.

